# Presaccadic attentional shifts are not modulated by saccade amplitude

**DOI:** 10.1101/2024.11.05.622100

**Authors:** Luan Zimmermann Bortoluzzi, Estêvão Carlos-Lima, Gabriela Mueller de Melo, Melissa Hongjin Song Zhu, Gustavo Rohenkohl

## Abstract

Humans constantly explore the visual environment through saccades, bringing relevant visual stimuli to the center of the gaze. Before the eyes begin to move, visual attention is directed to the intended saccade target. As a consequence of this presaccadic shift of attention (PSA), visual perception is enhanced at the future gaze position. PSA has been investigated in a variety of saccade amplitudes, from microsaccades to locations that exceed the oculomotor range. Interestingly, recent studies have shown that PSA effects on visual perception are not equally distributed around the visual field. However, it remains unknown whether the magnitude of presaccadic perceptual enhancement varies with the amplitude of the saccades. Here, we measured contrast sensitivity thresholds during saccade planning in a two-alternative forced-choice (2AFC) discrimination task in human observers. Filtered pink noise (1/f) patches, presented at four eccentricities scaled in size according to the cortical magnification factor were used as visual targets. This method was adopted to mitigate well-known eccentricity effects on perception, thereby enabling us to explore the effects associated to saccade amplitudes. First, our results show that saccade preparation enhanced contrast sensitivity in all tested locations. Importantly, we found that this presaccadic perceptual enhancement was not modulated by the amplitude of the saccades. These findings suggest that presaccadic attention operates consistently across different saccade amplitudes, enhancing visual processing at intended gaze positions regardless of saccade size.

## INTRODUCTION

Many animals - including humans - employ eye movements to actively sample the visual environment^[76]^. In most cases, this scanning is carried out by saccadic eye movements that constantly and abruptly change the visual input to the brain. This poses a great challenge for visual processing, as most of the visual sensory areas are retinotopically organized. Perhaps not surprisingly, several perceptual effects have been observed around the onset of saccadic eye movements^[3;25]^. One of the most robust effects is the presaccadic shift of attention (PSA)^[41]^, observed by the enhancement of perceptual processing of saccade-targeted objects.

Since PSA was first observed, studies have described several effects of presaccadic attention on visual perception (for reviews, see^[9;41]^). For instance, PSA was shown to enhance both contrast sensitivity^[30]^ and spatial resolution^[38]^ in the periphery of the visual field. Similar effects have been observed in the neural response of many visual areas^[48]^. However, recent studies have shown that PSA effects are not equally distributed across the visual field^[30;31]^. PSA effects have been observed in a wide variety of stimuli eccentricities[7;20;30;38–40;42;54;58;64], from microsaccades^[26;66]^ (20 arcmin) to eccentricities that exceed the oculomotor range (45 dva)^[32]^. In the seminal work by Deubel and Schneider (1996)^[20]^, PSA was tested in three eccentricities (3.91, 5 and 6.09 dva). Even though the difference in eccentricity values was arguably too small to test for potential differences in PSA effects, there seems to be a reduction in discrimination performance with eccentricity (especially comparing the 3.91 dva versus 6.09 dva conditions). However, since in their study the visual targets had a fixed size (0.52 and 1.05 dva for width and height, respectively), overall performance decayed with eccentricity, hindering any attempt to interpret their results in terms of saccade amplitude.

Eccentricity effects on visual perception have also been investigated in conditions where attention is spatially allocated without concurrent eye movements - i.e., covert attention^[2;10;11;14;17;22;35;68]^. Similar to Deubel and Schneider’s results, visual performance tends to decrease for both endogenous (top-down) and exogenous (bottom-up) covert attention across eccentricities when stimuli sizes are fixed^[12;22;68]^. However, when stimuli sizes are scaled according to the cortical magnification factor (M-scaled), these results change. The effects of eccentricity on visual performance are strongly attenuated for endogenous - but not exogenous - attention^[12;17;68]^. Nevertheless, it is not possible to deduce the impact of eccentricity on PSA from studies of covert attention, as these processes appear to rely on distinct computations^[42]^.

Successful saccadic eye movements rely on a large network of brain areas^[49]^. Areas such as frontal eye fields (FEF) and superior colliculus (SC) are known for being involved in the planning of saccades^[8;43;75]^. These areas are topographically organized, and microstimulation of specific sites produces saccadic movements of fixed directions and amplitudes^[61;63]^. Correspondingly, the firing rates of motor and visuomotor cells in these areas increase prior to saccade execution to locations relative to the fixation position^[8;50]^. This pattern of activity is referred to as *movement fields*. Interestingly, FEF and SC are interconnected with several visual sensory areas^[18;45;60;67]^. Microstimulation and pharmacological manipulations of FEF induced attention-like effects in visual areas (V4) and behavior^[46;47;52]^. A recent study has shown that transcranial magnetic stimulation (TMS) of FEF during saccade preparation modulates PSA effects^[29]^. For these reasons, both areas have been associated with PSA^[41;48]^.

Given the close relationship between oculomotor planning and visual perception, here we asked whether presaccadic attentional shifts are modulated by the size of the planned saccades. Since the oculomotor activity involved in the programming of saccades does not vary with different amplitudes, we expected that PSA effects would be uniform across eccentricities. Our results confirmed this prediction. We found that saccade preparation enhanced contrast sensitivity in all tested locations, and that this enhancement was not modulated by saccade amplitude.

## RESULTS

In this study, we investigated the effect of saccade amplitude on presaccadic attentional shifts. We developed a simple task to compare contrast sensitivity prior to saccades executed to targets presented at four eccentricities (Figure 1A-B). Because we were interested in the effect of saccade amplitude, target sizes were corrected using a magnification factor to minimize the effect of visual eccentricity. Sixteen subjects performed a two-alternative forced choice (2AFC) discrimination task, in which they had to report the orientation (clockwise or counterclockwise) of a filtered pink noise target presented at the saccade (towards) or the opposite location (away). Discrimination performance was measured online using an adaptive psychometric procedure (Best PEST), targeting the contrast level for 80% orientation discrimination accuracy. Figure 1C-F shows an example of the staircase procedure applied to one subject in all sessions.

**Figure 1:**
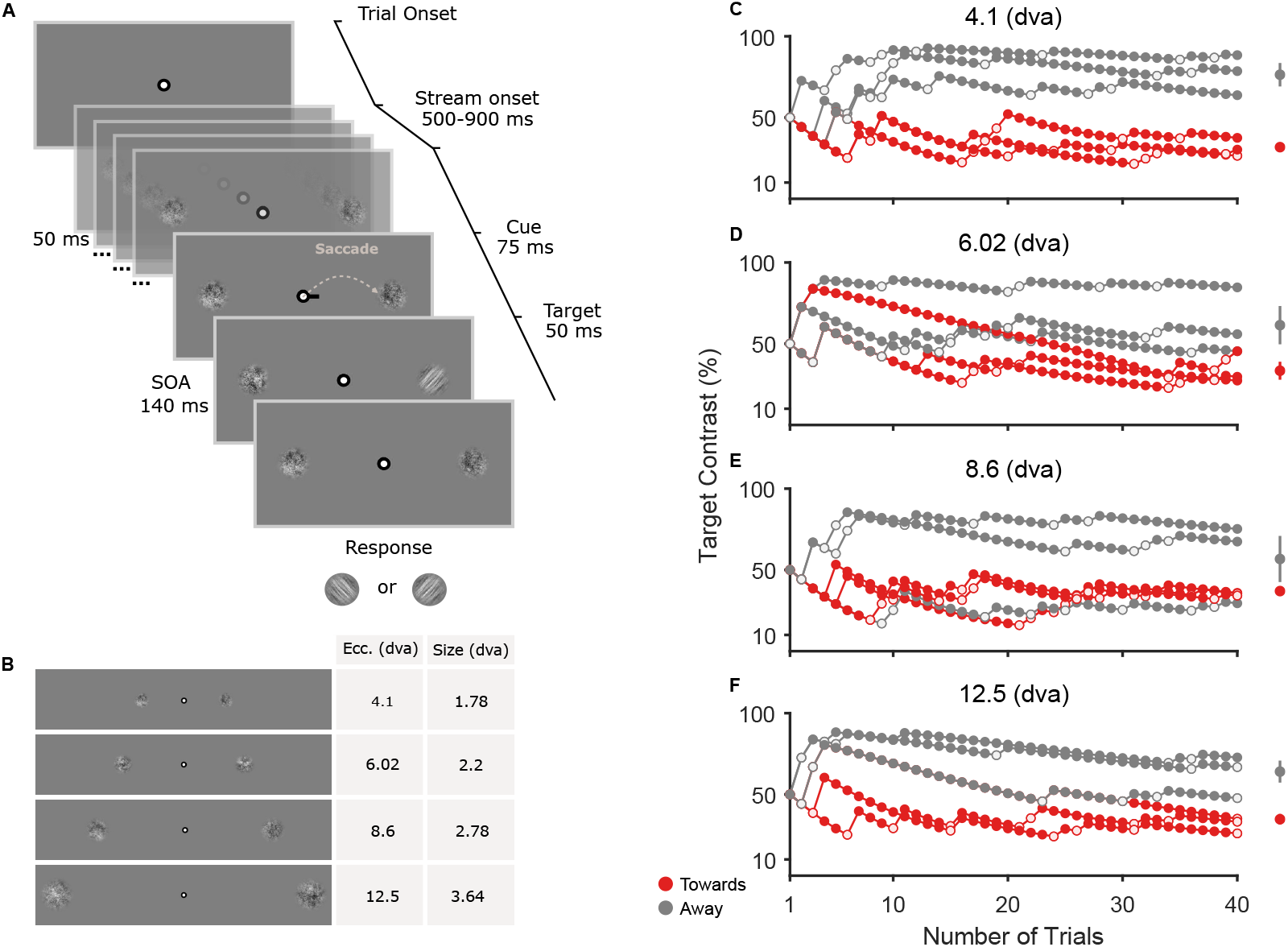
Task design and staircase result. (A) Trial example. Subjects were instructed to look at a fixation point (FP) for at least 500 ms at the beginning of each trial. Then, a stream of pink noise stimuli would be presented on both sides of the FP and remain on the screen until the end of the trial. After a random period of 500-900 ms, a saccadic cue (black line) would indicate the side where a saccade should be executed. After an SOA of 140 ms, a visual target (filtered pink noise) with a clockwise (CW) or counterclockwise (CCW) orientation would appear either on the left or right side of the screen for 50 ms. Subjects had to discriminate the orientation of the target at the end of the trial. (B) Stimuli eccentricities and sizes. (C-F) Example of the staircase procedure applied separately for each stimulus eccentricity. Target contrast is shown in function of the number of trials. Each staircase subplot shows the subject’s performance for cue towards (red) and away (gray) from the target. Filled and open dots represent correct and incorrect discrimination responses, respectively. On the right, the mean and standard error for each condition are shown. The mean was calculated using the means of the posterior distribution given by the staircase procedure.

### Saccade amplitude does not affect presaccadic attention

To investigate the effect of saccade amplitude on contrast sensitivity, we performed a two-way repeated measures ANOVA on individual threshold values. As expected, this analysis showed that contrast thresholds were lower when saccades were planned towards the target than in the away condition (Figure 2, F(1,15) = 81.94, p < .0001). Our analysis also revealed a main effect of stimulus eccentricity (F(3,45) = 9.339, p = .0005). Posthoc comparisons showed that targets presented at intermediary eccentricities led to lower contrast thresholds when compared to the nearest eccentricity (6.02-4.1 dva, p = .0001; and 8.6-4.1 dva, p = .001). Most importantly, we found no interaction between the cue direction (Towards and Away) with the target eccentricity condition (Figure 3A, F(3,45) = 0.03, p = .98). A linear regression analysis was performed to further examine the relationship between target eccentricity and presaccadic perceptual enhancement. To do so, we subtracted saccade-towards from -away condition for each stimulus eccentricity. This analysis confirmed the weak relationship between PSA and saccade amplitude (Figure 3B-C, slope: -0.417, p = .51469). These results showed that the presaccadic shifts of attention are not modulated by the amplitude of the saccade being prepared to the target location.

**Figure 2:**
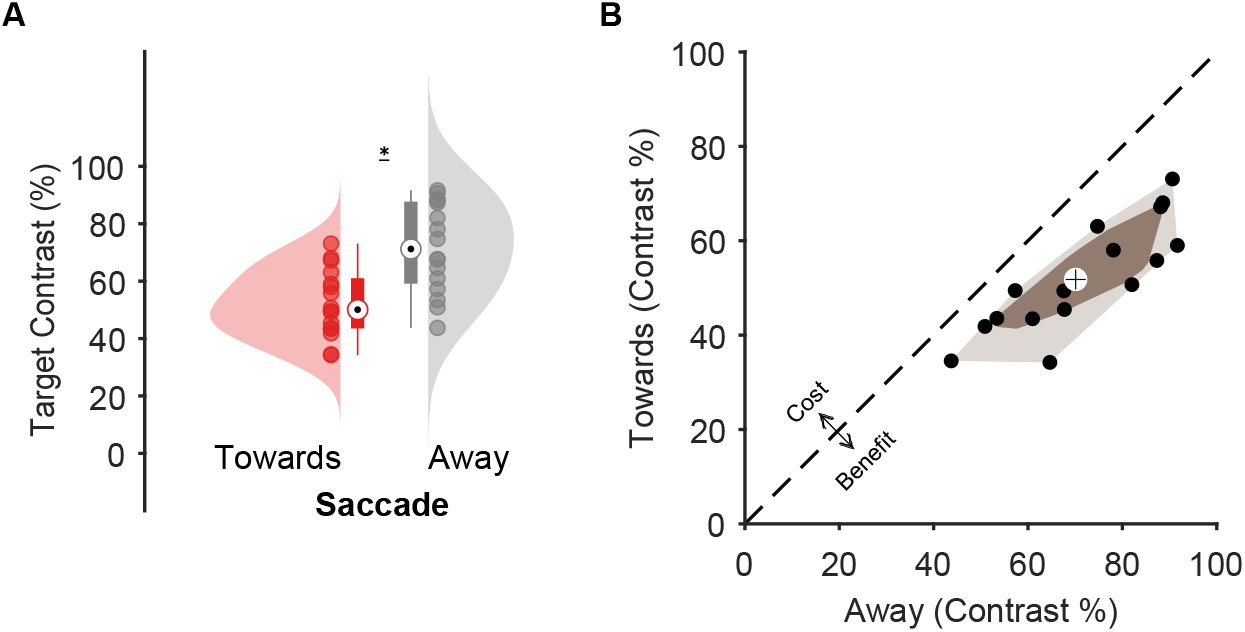
Effect of presaccadic attention on contrast sensitivity. (A) Target contrast as a function of cue direction: Towards and Away conditions are shown in red and gray, respectively. The black dot on each boxplot represents the median and the whiskers represent the upper and lower value. The colored clouds represent the density distribution given by the inside dots, which represent subjects’ performance individually. The asterisk indicates the statistical significance (p < .0001). (B) Bagplot for contrast sensitivity for towards vs away cue condition. The brown polygon (bag) contains 50% of the data (black dots) whereas the fence, in a lighter brown color, contains the remaining non-outlier data. The median is indicated by a cross.

**Figure 3:**
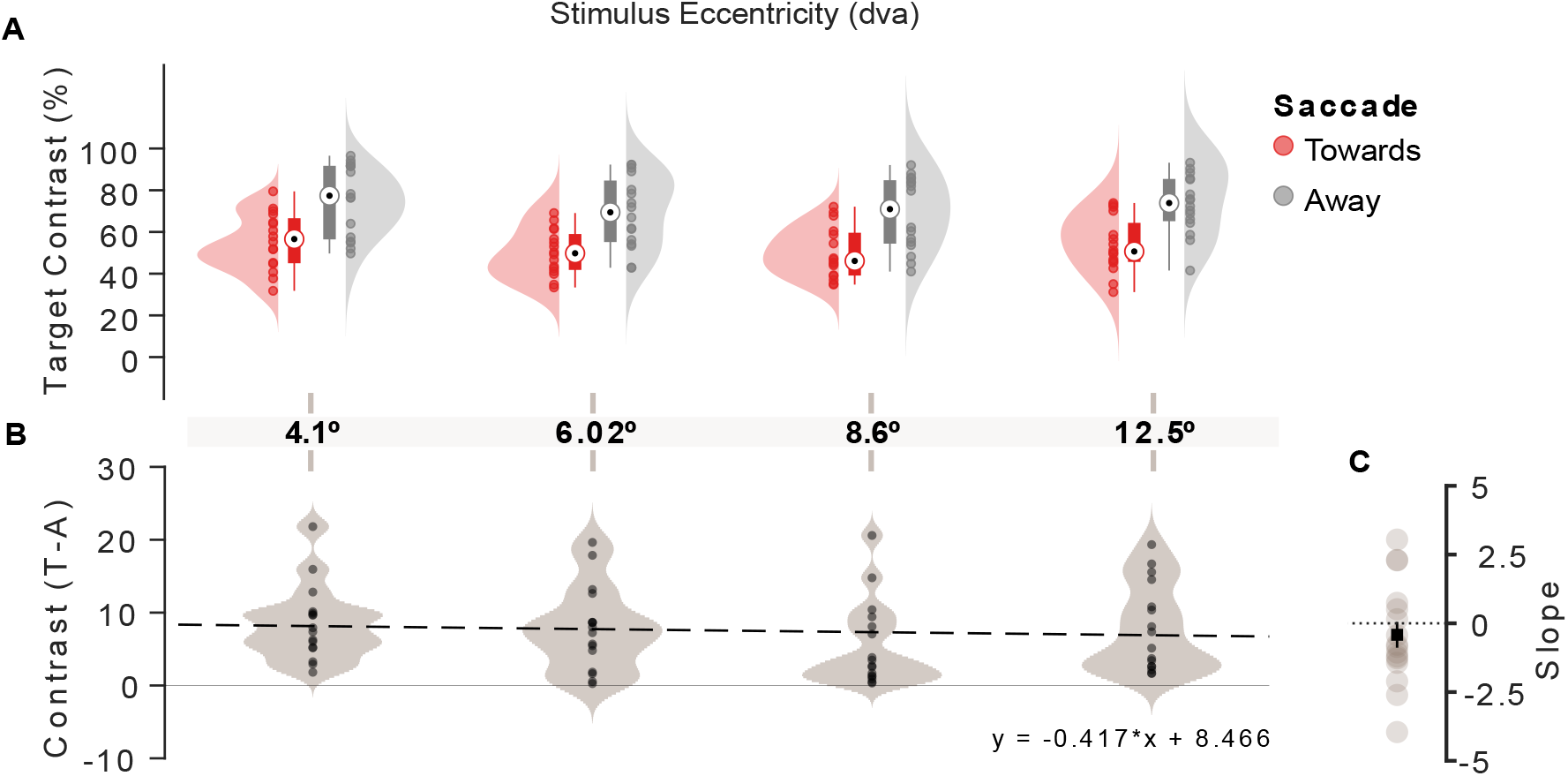
Presaccadic attention effects in all tested eccentricities. (A) Target contrast as a function of cue direction for each stimulus eccentricity: Towards and Away conditions are shown in red and gray, respectively. The black dot on each boxplot represents the median and the whiskers represent the upper and lower value. The colored clouds represent the density distribution given by the inside dots, which represent subjects’ performance individually. (B) Linear regression of the data and its violin plot distribution. (C) Slope of each subject’s linear regression fit. Mean and s.e. are indicated by the black square.

### Control analyses - saccade parameters

One possible confound in our study could be raised if saccades had different latencies for the different eccentricities. It is known that PSA effects are strongest when the target is presented just before saccade onset^[58]^. Because we used a fixed cue-target interval, if PSA effects decayed with eccentricity, but saccade latency increased, for example, then one effect could be canceling the other (i.e., targets in the nearest eccentricities would be presented closer to saccade onset). To evaluate this possible confound, we performed a two-way repeated measures ANOVA on saccadic reaction times (SRTs) across eccentricities. This analysis did not reveal a significant effect of cue direction on SRT (F(1,15) = 2.3797, p = .14375). There was a significant effect of Eccentricity on SRT (F(3,45) = 19.903, p < .0001). Post-hoc analyses showed that SRT tends to decrease with eccentricity (Figure 4A-D) [4.1-6.02 dva, permuted-p = .004; 4.1-8.6 dva, permutedp < .0001; 4.1-12.5 dva, permuted-p < .0001; 6.02-8.6 dva, permuted-p < .0001; 6.02-12.5 dva, permuted-p < .0001; 8.6-12.5 dva, permuted-p = .005]. First, it should be noted that even though they are significant different, the SRT differences between eccentricities are very small (3.60 ms for 4.1-6.02 dva; 9.21 ms for 4.1-8.6 dva; 12.31 ms for 4.1-12.5 dva; 5.60 ms for 6.02-8.6 dva; 8.70 ms for 6.02-12.5 dva; 3.09 ms for 8.6-12.5 dva). Most importantly, we found no interaction between the factors (F(3,45) = 1.381, p = .26486). If SRTs had affected our main results, we would not have seen the same pattern of results throughout stimulus eccentricities (see Figure 3), which was not the case.

**Figure 4:**
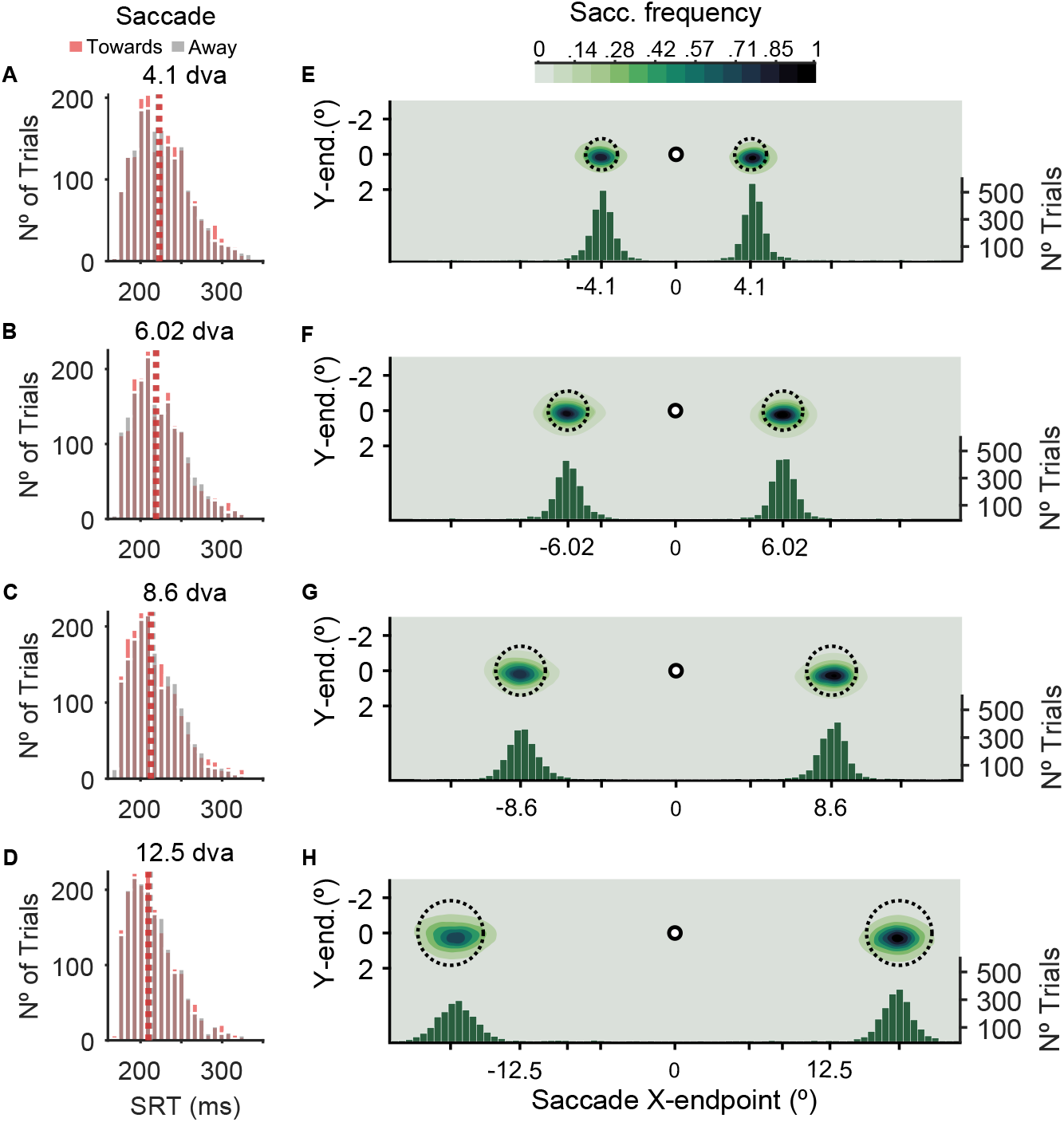
(A-D) Histograms depicting saccade reaction time (SRT) for cue direction towards (red) and away (gray) from the target for each eccentricity. Dashed lines represent the mean SRT. (E-H) Density distribution of saccade landing position for each stimulus eccentricity for all subjects. Dashed circles represent noise stimuli. Left Y-axis: vertical landing position; X-axis: horizontal landing position. Right Y-axis: Histogram depicting the number of trials as a function of saccade landing position for each stimulus eccentricity.

Another concern could be related to saccade accuracy. Even though it is known that PSA is associated with saccade planning and not execution^[74]^, and that it occurs even when saccade endpoints fall as far as 1.5 degrees away from the edge of the target stimulus^[32]^, one could still argue that highly inaccurate saccades could affect our main results. We performed an analysis to investigate the effect eccentricity on saccade accuracy. Saccade accuracy data were corrected for each eccentricity according to the cortical magnification factor (see Methods). This analysis revealed that saccade accuracy was equivalent for both cue directions (F(1,15) = 1.3491, p = .26) and all eccentricities (F(3,45) = 0.22468, p = .73) (Figure 4E-H). These results rule out the possibility of any saccade parameters biasing the PSA effects in our experiment.

## DISCUSSION

Presaccadic shifts of attention have been observed using a wide range of stimulus eccentricities[7;20;30;38–40;42;54;58;64]. Given the intrinsic link between stimuli eccentricities and saccade amplitudes, we expected that visual attention could be similarly engaged for saccades of different sizes. However, until now, the evidence for this assumption could only be obtained by combining results from several independent studies. Therefore, here we directly investigated if presaccadic attentional effects on visual perception are modulated by saccade amplitudes. We employed filtered pink noise (1/f) patches as target stimuli, scaled in size according to the cortical magnification factor. This approach aimed to mitigate the well-known effects of stimulus eccentricity on contrast sensitivity, allowing us to examine the effects of saccade amplitude on presaccadic shift of attention (PSA). Our results indicated that saccade preparation enhanced contrast sensitivity in all tested locations. Furthermore, we showed that this presaccadic perceptual enhancement was not modulated by the amplitude of the saccade. Control analyses suggested that this result could not be explained by differences in saccade statistics.

It has long been established that oculomotor areas, such as the frontal eye fields (FEF) and superior colliculus (SC), are involved in attentional selection^[5;6]^. Unsurprisingly, several studies have suggested that PSA also depends on the involvement of these areas^[4;46;47;51]^. Stimulation of specific sites in FEF and SC will produce saccades with a fixed direction and amplitude relative to the current eye position^[61;63]^. Crucially, these saccade vectors are defined by the site that is stimulated and not by the intensity of the stimulation^[8;53;62]^. Therefore, the lack of interaction between saccade amplitude and presaccadic enhancement observed in our study aligns with the functional organization of these brain areas.

It is well known that visual perception is not equally distributed across the visual field (for a review, see^[70]^). Instead, contrast sensitivity is much reduced in the upper vertical meridian^[1;36;59]^. Interestingly, recent studies have shown that PSA effects are unevenly allocated across the visual space^[30;31]^. These studies reveal a polar angle asymmetry, in which the contrast sensitivity enhancement before upward vertical saccades is highly diminished - or even extinguished. We did not find any asymmetry in the magnitude of contrast enhancement before horizontal saccades for stimuli at different eccentricities.

One interesting takeaway from our study is that PSA effects are probably not linked to motor training. Studies on the ecology of saccades have shown that the natural distribution of human saccade amplitudes is far from uniform^[23;34]^. We execute nearly twice as many 5 degree saccades than 10 degree ones in our daily activities^[23]^. If PSA was associated with the frequency of saccade amplitudes, then we would expect that our results followed the natural distribution. Instead, our results showed that PSA effects - at least for the eccentricities tested - seem to be uniformly distributed.

Evidence from psychophysical studies suggests that PSA is determined by the intention of the saccade goal and not the execution accuracy^[20;28;32;74]^. Given that saccades do not always land at the intended location - often undershooting the target - a large number of studies, starting from Deubel and Schneider’s (1996)^[20]^ seminal work, have proposed that in spite of these saccade errors, visual perception is always enhanced at the intended location. Wollenberg and colleagues (2018)^[74]^ tested perceptual enhancement prior to averaging saccades (i.e., when saccades land in between two closely located saccade targets), which allowed the dissociation of visual sensitivity modulations at the intended saccade goal versus at the endpoint of the saccade. Their results showed that, while accurate saccades toward one of the targets were associated with presaccadic enhancement, there was no attentional facilitation at the saccade endpoint of an averaging saccade. Another evidence comes from a study by Hanning and colleagues (2019)^[32]^ showing PSA effects to targets located outside the oculomotor range, even though, in these conditions, saccades always fell short of the target location. Our results agree with these proposals. One would expect that if PSA was related to the execution - and not intention - of the saccade, perceptual enhancement effects would decrease with saccade amplitude. Conversely, our results show that the contrast sensitivity enhancement was equal prior to saccades ranging from 4.1 to 12.5 dva.

The majority of the evidence on the relationship between visual attention and target eccentricity comes from visual search studies^[10;11;13;15;16;73]^. These studies have shown that performance consistently decreases with the eccentricity of targets in a search array. However, this eccentricity effect seems to diminish when correcting the array stimuli for cortical magnification (^[12;15]^, cf.^[68]^), or when a pre-cue is presented indicating the target location^[16]^. More recently, using a classic Posner-like (covert) attentional task, Jigo and Carrasco (2020)^[35]^ have shown that a central cue is capable of enhancing the contrast sensitivity of targets presented at different eccentricities (0, 3, 6, and 12 dva), and across a wide range of spatial frequencies. Similarly, Hanning and Deubel^[27]^ have shown that visual performance is equated for 1/f stimuli when a pre-cue is presented.

It is well known that humans perceive the visual world in a stable, continuous way across saccades. This visual stability is thought to be due (but not only) to trans-saccadic integration. This phenomenon can be understood as the fusion of the peripheral low-resolution information during the programming of a saccade with the foveal high-resolution information after the execution of a saccade. Trans-saccadic integration contributes to the assimilation of several different stimulus parameters, such as color^[71]^, form^[19]^, orientation^[72]^, information about stimulus location^[56]^, and so on. Although PSA was often thought to be a possible contributing mechanism to this integration, it was not until recently that studies have found supporting evidence^[65;69]^. If one considers that PSA plays an important role in trans-saccadic integration (and thus, contributes to visual stability across saccades), it is reasonable to assume that these shifts of attention would occur before any saccade, irrespective of their amplitude. Therefore, our main results contribute, even though indirectly, to the trans-saccadic integration hypothesis.

Our work raises several interesting questions to be investigated in future studies. Here, we used a fixed cue-target SOA of 140 ms. This prevented us from investigating the time course of the perceptual enhancement before saccade onset. It is possible that differences in PSA for different saccade amplitudes might emerge when analyzing its temporal profile more carefully. In our study, we also limited the analysis of PSA effects to stimuli up to 12.5^*°*^ in eccentricity. This was done for two main reasons. First, we wanted to avoid presenting stimuli near the blind spot, found around 15^*°*^ in the temporal horizontal hemispace for each eye. Second, under natural, free-viewing conditions, head movements begin to contribute to gaze shifts for locations larger than 20^*°*^ in both humans and non-human primates (for a review, see^[24]^). It is possible that PSA effects are attenuated for saccades larger than 20^*°*^, when head movements start being engaged. Finally, we have limited stimulus position to the horizontal meridian, as horizontal saccades are more common than vertical ones under natural behavior conditions^[23]^, and because - as described above - there are known differences in performance across the visual field^[30;31]^.

In conclusion, our study showed that PSA enhances contrast sensitivity equally before saccades of up to 12.5^*°*^. Moreover, our results suggest that any saccade amplitude within the range used here can be applied to future PSA studies without impairments to the phenomenon.

## MATERIAL AND METHODS

### Participants

Sixteen subjects participated in this study (10 females and 6 males; mean age 26.75). Except for three authors (L.Z.B, E.C.L, M.H.S.Z), all participants were naive to the purposes of the study. All participants had normal or corrected-to-normal vision, with 11 of them having dominance in the right eye. Before participating in any part of the study, the subjects read and signed a consent form approved by the local Ethics Research Committee (approval number: 25333219.5.0000.5464).

### Apparatus

The experiment was carried out in a quiet, dimly lit room, where the participants sat comfortably in front of a computer screen and positioned their heads on a chin-rest. Stimuli were presented at a distance of 57 cm from the subjects’ eyes on a high-resolution screen (1920 × 1080, refresh rate 120 Hz) controlled using the Psychtoolbox toolbox^[37]^ in MATLAB^[33]^. Manual responses were recorded using a response box (ResponsePixx). Eye position was monitored at 1KHz sampling rate using the desktop mount EyeLink Plus 1000 (Eye Link Plus 1000 SR Research).

### Stimuli and Task

Each trial started with a fixation point on a gray background. The fixation point consisted of a white dot with a diameter of 0.3 dva, within a black circle with a diameter of 0.64 dva. After 500 ms of central fixation - within a 2.35 dva radius window - two visual streams (50 ms SOA) of pink noise stimuli were presented at the left and right sides of the fixation dot until the end of the trial. After a random interval between 500-900 ms, a central line (0.41 dva in length and 0.14 dva in width) pointed to the left or right side (75 ms duration), signaling a spatial cue. Subjects were instructed to execute a saccade as fast as possible to the stimuli location at the cued side. Then, following an interval of 65 ms (140 ms cue-target SOA), an orientation-filtered pink noise patch (40^*°*^ or -40^*°*^ relative to vertical) was presented at the cued or uncued location for 50 ms. Subjects were instructed to discriminate the target’s orientation (clockwise or counterclockwise) by pressing a button on the response box. The saccadic cue was not predictive of the future target location (i.e., 50% validity).

Visual performance was tested at four different logarithmically spaced eccentricities (4.1, 6.02, 8.6, and 12.5 dva), and a magnification factor (Mscaling) was applied to the stimuli sizes (1.78, 2.2, 2.78, and 3.64 dva) (Figure 1B). Using the magnification factor in combination with the pink noise stimuli ensured that visual performance would remain similar across all eccentricities^[12;27;36]^.

### Procedures

Participants performed three sessions of 320 trials (960 trials in total). Each session consisted of 20 short blocks of 16 trials, with approximately 30-40 minutes of duration. At the beginning of each block, a message displayed on the screen indicated the stimulus eccentricity in that block.

The eccentricity was randomized between blocks. Within the same block, trials were repeated if: the subjects’ eyes deviated from the fixation point before cue onset; saccade reaction times were faster than 190 ms or slower than 350 ms; saccades were executed to an uncued location; no saccade was detected during the whole trial. Subjects received visual feedback indicating unsuccessful saccade execution at the end of these trials (the fixation point turned yellow). Subjects were allowed to rest between blocks.

Before starting each experimental session, the participants performed a short training session similar to the main experiment. During training, in addition to the saccade feedback, participants also received a second feedback regarding the discrimination performance (the fixation point turned green for correct or red for incorrect responses). In the first experimental session, participants received detailed instructions before the training session.

### Staircase Procedure

A Best Pest staircase procedure^[55]^, was used to measure the contrast level for 80% correct discrimination responses in each condition, using Palamedes toolbox^[57]^. For each session, eight staircases were applied concurrently to derive the targets’ contrast level for each of the eight experimental conditions: Cue (Towards and Away) and Stimulus Eccentricity (4.1, 6.02, 8.6, and 12.5 dva). Each staircase condition consisted of 40 trials. The contrast threshold for each condition was derived from the mean of the posterior distribution. Then, before calculating the overall mean across all sessions for each condition, we excluded outlier data (1.56% of all staircase procedures) for which the mean of a specific staircase condition was higher than 0.5 log compared to the other two means in the same condition. Moreover, we also excluded the data from an entire session of a subject who initially misunderstood the instructions and took three times longer than usual to complete the session. Running the same analysis but without excluding the outlier values yielded similar results than with exclusions.

### Data Analysis

The analysis of discrimination performance was based on the contrast level needed to achieve 80% correct discrimination in each condition (see staircase procedure above). In addition to discrimination thresholds, we also considered as dependent variables saccade reaction times (SRT) and saccade accuracy. Two-way repeated-measures ANOVAs were implemented for all dependent variables. In the results section, p-values are shown with Greenhouse-Geisser correction due to sphericity violation. Bonferroni correction was applied to post-hoc analysis on discrimination performance. Since SRT and saccade accuracy did not show normal distribution, according to the Kolmogorov-Smirnov test, post-hoc analyses were run using a non-parametric permutation test^[44]^. Given that the stimuli sizes were corrected by the magnification factor, saccade accuracy data was corrected for each eccentricity by dividing the post-saccade position by the stimulus radius.

### Eye movements

In addition to the ‘online’ eye movement analysis, we used a velocity-based saccade detection algorithm to detect saccades offline, which provides more detailed information regarding saccade parameters^[21]^. Saccades onset and offset were detected based on the current eye velocity at the point in which median eye velocity exceeded 5 SDs. This analysis was used only for SRT and saccade accuracy data. A total of 497 (3,24%) trials were excluded.

## ACKNOWLEDGMENTS

This work was supported by the São Paulo Research Foundation (FAPESP) and by the Institute of Biosciences - IB/USP. We thank everyone who volunteered to participate in our experiment. We would also like to thank all our lab colleagues for all the suggestions given throughout the development of the study.

## AUTHOR CONTRIBUTIONS

Conceptualization and methodology: G.R., L.Z.B., E.C.L., M.H.S.Z., G.M.M.

Research: G.R., L.Z.B., E.C.L.

Software: L.Z.B., G.M.M.

Data collection: L.Z.B., M.H.S.Z.

Data analysis: L.Z.B., E.C.L., G.R.

Visualization: L.Z.B.

Writing: L.Z.B., G.R.

Manuscript Review: G.R., L.Z.B., E.C.L., G.M.M.

Funding acquisition: G.R.

## DATA AVAILABILITY

The datasets generated in this study can be acquired from the corresponding author on request.

